# What is learned determines how Pavlovian conditioned fear is consolidated in the brain

**DOI:** 10.1101/2022.12.05.519226

**Authors:** Jessica Leake, Dana M. Leidl, Belinda P. P. Lay, Justine P. Fam, Madeleine C. Giles, Omar A. Qureshi, R. Frederick Westbrook, Nathan M. Holmes

## Abstract

Activity in the basolateral amygdala complex (BLA) is needed to encode fears acquired through contact with both innate sources of danger (i.e., things that are painful) and learned sources of danger (e.g., being threatened with a gun). However, within the BLA, the molecular processes required to consolidate the two types of fear are not the same: protein synthesis is needed to consolidate the first type of fear (so-called first-order fear) but not the latter (so-called second-order fear). The present study examined why first- and second-order fears differ in this respect. To do so, it used a range of conditioning protocols in rats and assessed the effects of a BLA infusion of the protein synthesis inhibitor, cycloheximide, on fear to first- and second-order conditioned stimuli. The results revealed that the differential protein synthesis requirements for consolidation of first- and second-order fears reflect differences in what is learned in each case. Protein synthesis in the BLA is needed to consolidate fears that result from encoding of relations between stimuli in the environment (stimulus-stimulus associations, typical for first-order fear) but is not needed to consolidate fears that form when environmental stimuli associate directly with fear responses emitted by the animal (stimulus-response associations, typical for second-order fear). Thus, the substrates of Pavlovian fear conditioning in the BLA depend on the way that the environment impinges upon the animal. This is discussed with respect to theories of amygdala function in Pavlovian fear conditioning, and ways in which stimulus-response associations might be consolidated in the brain.

## Introduction

Some of our fears originate in experiences with innate sources of danger, for example, the fear produced by pain from burning embers and suffocation from smoke when caught in a bush fire. Other fears originate in our experiences with learned sources of danger, for example, being frightened by someone who is pointing a gun at you and demanding money is contingent on you having learned that guns are dangerous. Each of these experiences create enduring memories which can be retrieved and expressed in fear responses that prepare the body for fight or flight: for example, the smell of smoke when walking in the bush or the sight of the ATM where you were robbed at gun point.

Are fears acquired through contact with innate sources of danger the same or different to those acquired through contact with learned sources of danger? In the laboratory, the two types of fear can be studied using a protocol in which rats are first exposed to pairings of a novel stimulus (S1; e.g., a light) and brief-but-painful foot shock (unconditioned stimulus, US), and 48 hours later, to presentations of a second novel stimulus (S2; e.g., a tone) followed by the previously conditioned S1 (S2àS1). During subsequent test sessions, rats exhibit defensive or fear responses (e.g., freezing, potentiated startle) when re-exposed to either the S1 or S2. Importantly, fear of the S2 is not due to generalized fear from the S1, as controls exposed to explicitly unpaired presentations of the stimuli in training do not exhibit this fear (Rizley and Rescorla, 1972; Barnet et al., 1991; Parkes and Westbrook, 2010; Witnauer and Miller, 2011; Holmes et al., 2013). Instead, fear of the S2 is the result of learning that occurs when this stimulus is paired with the already-conditioned S1. Pavlov called this learning second-order conditioning to distinguish it from the learning that results from S1-shock pairings, which he called first-order conditioning (Pavlov, 1927).

First- and second-order conditioned fears both require activity in the basolateral amygdala complex (BLA) for their acquisition and retrieval/expression; and both are expressed in a common set of defensive responses (e.g., freezing, potentiated startle; Gewirtz & Davis, 1997, 2000; Holmes et al., 2013). However, the two types of fear differ in important ways. One difference relates to the processes required for their consolidation in long-term memory. For example, consolidation of first-order fear requires synthesis of new proteins in the BLA (Schafe and LeDoux, 2000; Maren et al., 2003; Desgranges et al., 2008; Kochli et al., 2015). These proteins support physical/structural changes (e.g., changes in the size, shape and number of dendritic spines) that allow subsequent presentations of the CS to more effectively activate neurons in the BLA and, thereby, intra-amygdala circuits that coordinate defensive responses (Ostroff et al., 2010; Johansen et al., 2011). By contrast, consolidation of second-order fear occurs independently of protein synthesis in the BLA (Lay et al., 2018; Leidl et al., 2018). Rats that receive a BLA infusion of a protein synthesis inhibitor (e.g., cycloheximide) immediately after the session containing pairings of S2 and the already-conditioned S1 freeze just as much to S2 at test as vehicle-infused controls. Such findings have been taken to suggest that consolidation of fear to S2 exploits cellular changes that occur during the prior acquisition of fear to S1: hence, it is does not require *de novo* protein synthesis in the BLA (Leidl et al., 2018). However, at present, this hypothesis remains to be tested.

First- and second-order fears also differ with respect to what is learned in conditioning. This has been demonstrated in a series of studies which show that, subsequent to second-order conditioning: 1) habituating animals to the US undermines the expression of fear to S1 without affecting the expression of fear to S2; and 2) repeated S1 alone exposures extinguish its ability to elicit fear responses without affecting the expression of fear to S2. (Rizley and Rescorla, 1972; Rescorla, 1973a; Holmes et al., 2014). These findings have led to the proposal that while first-order conditioning is supported by an association between the sensory properties of the conditioned (S1) and unconditioned stimuli (i.e., a stimulus-stimulus association, S-S), second-order conditioning (as it is standardly conducted) is not. Instead, second-order conditioning is supported by an association between the S2 and fear responses triggered by the already-conditioned S1 (i.e., a stimulus-response association, S-R). Thus, an alternative explanation for the finding that protein synthesis is required to consolidate first- but not second-order fear is that the protein synthesis requirement is specific to a particular type of learning. That is, protein synthesis in the BLA is needed to consolidate S-S associations of the sort that typically form in first-order conditioning but is not needed to consolidate S-R associations of the sort that typically form in second-order conditioning.

The present study examined why protein synthesis is required to consolidate first- but not second-order fear in the BLA. It tested two hypotheses. The first is that second-order fear is consolidated via proteins that are synthesized during acquisition of first-order fear: hence, no further protein synthesis is required. To test this hypothesis, we shortened the interval between first and second-order conditioning from 48 h (used by Lay et al., 2018, Leidl et al., 2018) to just 10 min and examined the effects of a BLA cycloheximide infusion on consolidation of fear to the S1 and S2. Under these circumstances, consolidation of second-order fear could not be mediated by proteins synthesised during acquisition of first-order fear, as these would also be disrupted by the BLA infusion of cycloheximide. Thus, if the hypothesis is correct, we expected that the BLA cycloheximide infusion would disrupt fear to both the S1 and S2 at test.

The second hypothesis tested is that the protein synthesis requirement for consolidation of Pavlovian conditioned fear tracks the type of association produced by conditioning. We tested this hypothesis by examining the effect of a BLA cycloheximide infusion on consolidation of fear to S1 and S2 in two different second-order conditioning protocols. The first protocol was the one that is standardly used by our lab and others: in this protocol, each S2 presentation is immediately followed by a presentation of the already-conditioned S1 (i.e., the stimuli are presented *serially,* S2àS1), and S2 directly associates with fear responses elicited by the S1 (S2-fear) (Rizley and Rescorla, 1972; Holmes et al., 2014). The second protocol was one in which each S2 presentation perfectly overlaps with a presentation of the already-conditioned S1 (i.e., the stimuli are presented *simultaneously,* S2S1), and S2 comes to elicit fear indirectly via its association with the already-conditioned S1 (S2-S1, S1-shock; Rescorla, 1982). If protein synthesis in the BLA is needed to consolidate S-S associations of the sort that typically form in first-order conditioning but is not needed to consolidate S-R associations of the sort that typically form in second-order conditioning, we expected that the BLA infusion of cycloheximide would have different effects on fear to S2 in the two conditioning protocols. Specifically, we expected that the cycloheximide infusion would have no effect on fear to S2 in the standard protocol, where that fear is supported by a direct S2-fear association; but would disrupt fear to S2 in the modified protocol, where that fear is supported by integration of S2-S1 and S1-shock associations.

## Results

### Experiment 1

The first experiment sought to establish second-order conditioning using a truncated protocol in which S1-shock pairings and S2àS1 pairings were separated by ten minutes rather than two days (as was used previously, see Lay et al., 2018, Leidl et al. 2018). We aimed to demonstrate that rats could acquire fear to the S2 in the truncated protocol and that this fear is associatively mediated: due to the pairing of the S2 and S1 rather than generalization from the conditioned S1 or any intrinsic ability of S1 to condition fear (see Rizley & Rescorla, 1972). The design included three groups of rats. Rats in Group Paired-Paired (PP) were exposed to S1-shock pairings in stage 1 and S2àS1 pairings in stage 2. Rats in Group Paired-Unpaired (PU) were exposed to S1-shock pairings in stage 1 and explicitly unpaired presentations of S2 and S1 in stage 2. Rats in Group Unpaired-Paired (UP) were exposed to explicitly unpaired presentations of S1 and shock in stage 1 and S2àS1 pairings in stage 2.

Figure 1B shows levels of freezing during S1 presentations in stage 1. Averaged across groups, there was a significant increase in freezing across the four trials (*F*_(1, 20)_ = 26.52, *p* < 0.001). Unexpectedly, there was no significant difference in freezing levels between rats that received paired presentations of S1 and shock (groups PP and PU) and rats that received unpaired presentations (group UP) (*F*_(1, 20)_ = 0.15, *p* = 0.70). Further, there were no significant group x trend interactions (*F*_s_ < 2.8, *p* > 0.05). The failure to detect a difference in freezing between the paired and unpaired groups likely reflects context conditioning by the shock in the unpaired group.

**Figure 1.**
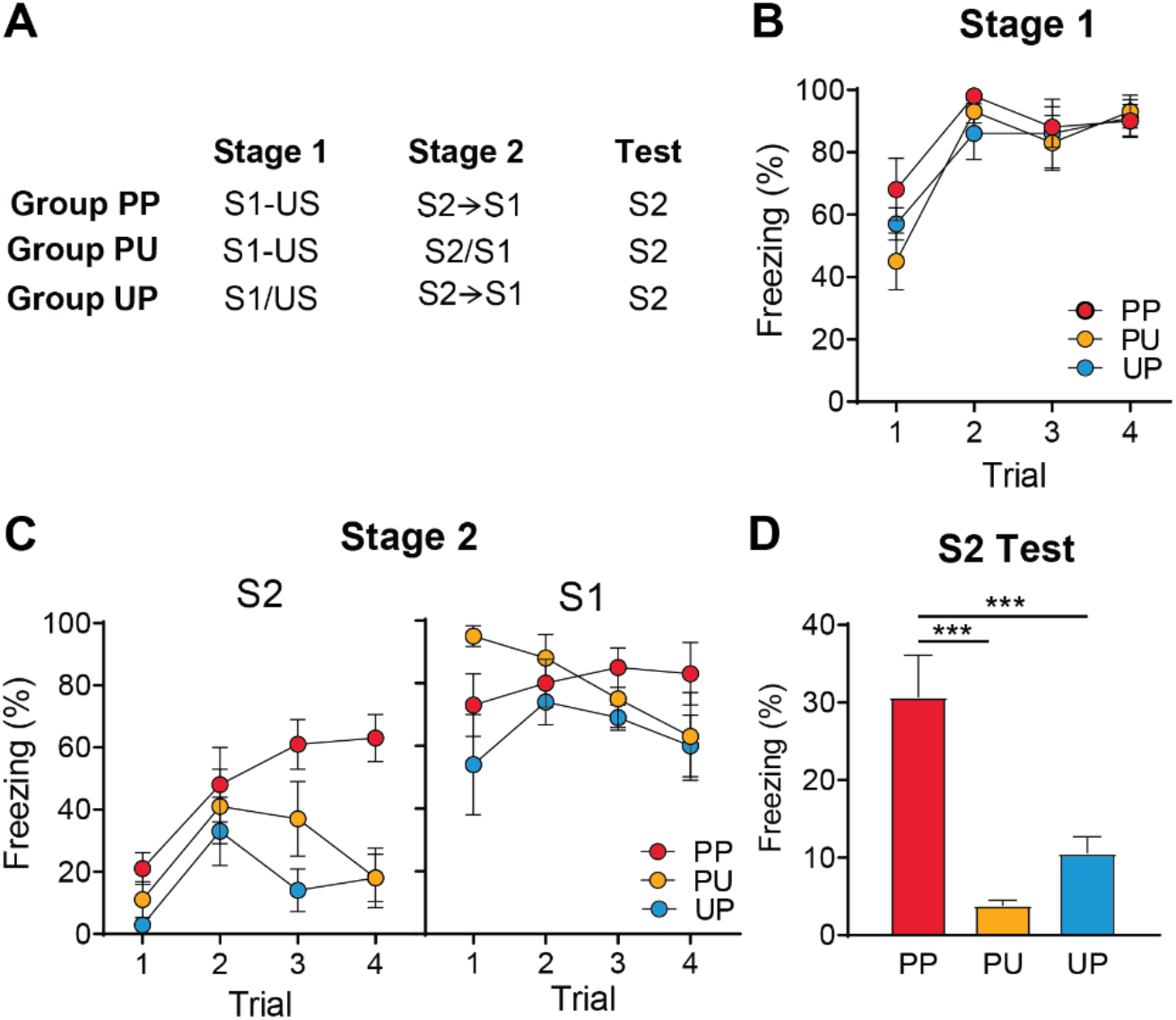
Freezing to S2 in the truncated protocol is due to second-order conditioning. (A) Experimental groups for conditioning and testing. A (à) indicates that the stimuli were presented in serial compound, whereas (/) indicates that the stimuli were explicitly unpaired (B) Mean (±SEM) levels of freezing to presentations of the S1 during stage 1. (C) Mean (±SEM) levels of freezing to presentations of the S2 and the S1 during stage 2. (D) Mean (+SEM) levels of freezing across test presentations of the S2.

Figure 1C shows levels of freezing during S2 presentations in stage 2. Averaged across groups, there was a significant increase in freezing to S2 across the four trials (*F*_(1,20)_ = 12.53, *p* <.005), indicating that rats acquired freezing to S2 across the session. Averaged across trials, rats in Group PP froze more to S2 that rats in Groups PU and UP (*F*_(1,20)_ =14.10, *p* < 0.005). Likewise, there was a difference in the rate of acquisition between rats in Group PP and those in Groups PU and UP (*F*_(1,20)_ = 10.40, *p* < 0.005). During stage 2, freezing to the S1 remained stable across the four trials (*F* < 1.0, p > 0.05). Averaged across trials, rats in Group UP froze less than rats in Groups PP and PU (*F*_(1,20)_ = 4.74, p <.05), thus confirming that responding to S1 in the latter two groups was due to its pairings with shock in stage 1.

At S2 test (Figure 1D), rats in Group PP froze significantly more than those in the other two groups (PP vs PU, *F*_(1,20)_ = 31.828, *p* < 0.0001, PP vs UP, *F*_(1,20)_ = 16.68, p< 0.0001). There was no significant difference in freezing between Groups PU and UP (*F*_(1,20)_ =1.87, *p* = 0.19). Together, these results indicate that freezing to S2 in the truncated protocol is conditional on its pairing with the conditioned S1 in stage 2, as well as the prior pairing of S1 with shock in stage 1. That is, this freezing to S2 is the result of second-order conditioning rather than generalization from the conditioned S1, or any unconditioned ability of S1 to condition freezing.

### Experiments 2a-c

The next series of experiments used the truncated protocol to test the hypothesis that proteins synthesized in the BLA during (or after) first-order conditioning are used to consolidate second-order conditioning. In each experiment, rats received S1-shock pairings in stage 1, followed ten minutes later by S2àS1 pairings in stage 2. The experiments differed with respect to the timing of the intra-BLA CHX infusion. Rats were infused with CHX either after S2àS1 pairings in stage 2 (Experiment 2a), after S1-shock pairings in stage 1 (Experiment 2b) or prior to S1-shock pairings in stage 1 (Experiment 2c).

The results of Experiments 2a-c are presented in Figure 2. Across all experiments, rats acquired fear to S1 and S2 during first- and second-order conditioning trials. Analysis of freezing across S1-shock pairings revealed a significant linear trend (Figure 2B, *F*_(1, 46)_ = 119.70, *p* < 0.001, Figure 2D, *F*_(1, 19)_ = 69.96, *p* < 0.001, Figure 2G, *F*_(1, 23)_ = 97.89, *p* < 0.001). There was no significant difference between groups in the rate of increase in freezing to S1 or in the overall level of freezing to S1 (*F*s < 2.5, *p* > 0.05). Freezing to S2 increased across its pairings with the conditioned S1, as evidenced by a significant linear trend (Figure 2B, *F*_(1, 46)_ = 119.70, *p* < 0.001, Figure 2D, *F*_(1, 19)_ = 69.96, *p* < 0.001, Figure 2G, *F*_(1, 23)_ = 97.89, *p* < 0.001). There were no significant differences between groups in the rate of increase in freezing to S2 or overall level of freezing to S2 (*F*s < 3, *p* > 0.05).

**Figure 2.**
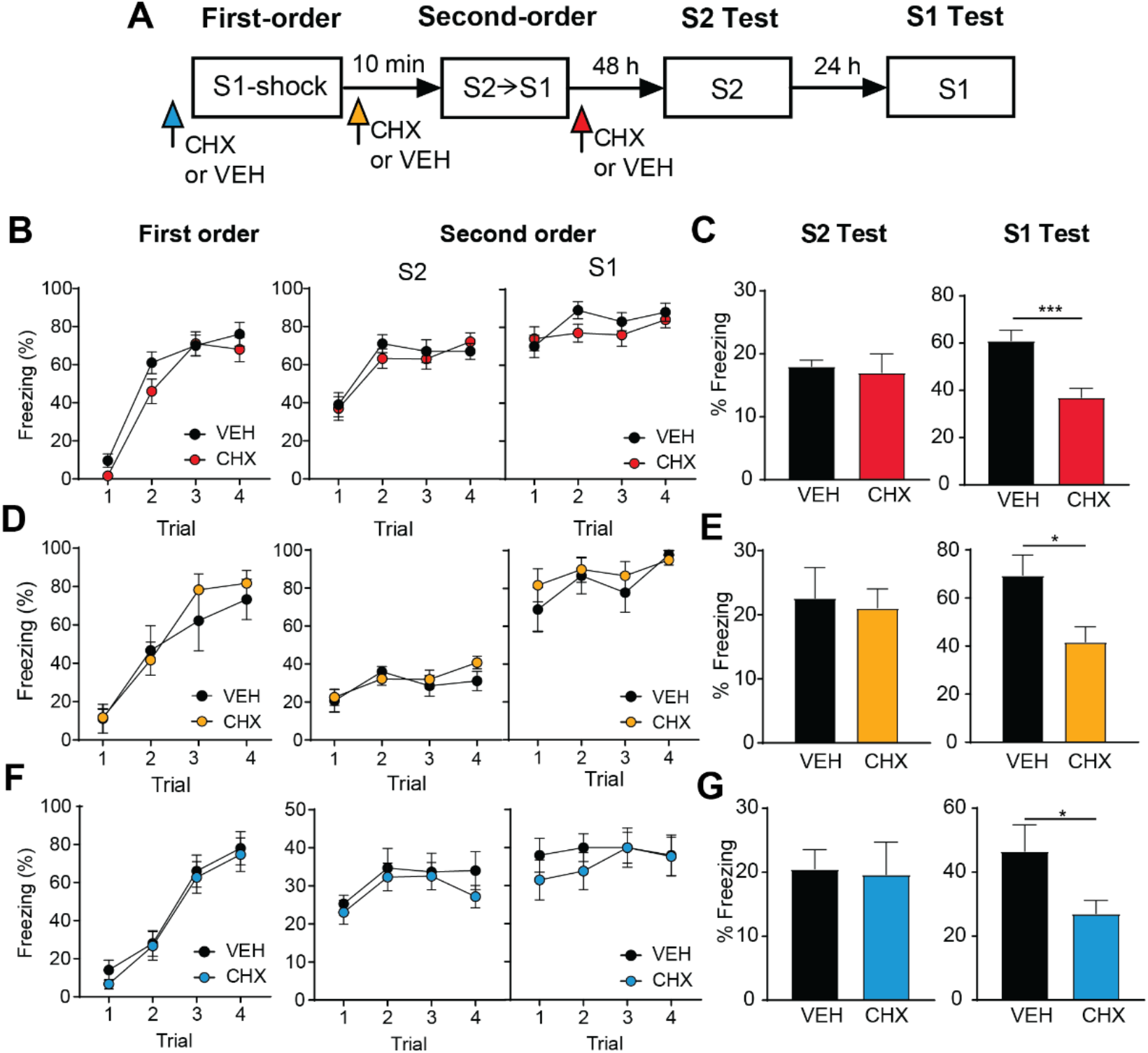
Protein synthesis in the BLA is required for consolidation of fear to S1 but not S2 in the truncated second-order conditioning protocol. (A) Experimental timeline of conditioning and testing for rats that received intra-BLA infusions of CHX or VEH. (B) Mean (±SEM) levels of freezing to presentations of the S1 during first-order conditioning. (C) Mean (±SEM) levels of freezing to presentations of the S2 and the S1 during second-order conditioning. (D-E) Mean (+SEM) levels of freezing across test presentations of the S2 (D) and the S1 (E).

In Experiments 2a and 2c, freezing to the S1 increased across S2àS1 shock pairings (Figure 2B, *F*_(1, 46)_ = 13.02,*p* < 0.001, Figure 2D, *F*_(1, 19)_ = 8.41, *p* < 0.001). Minimally, this indicates that four non-reinforced presentations of S1 was insufficient to produce any extinction. It could also indicate that the absence of shock had removed any shock-elicited escape attempts that would have interfered with freezing in stage 1. In Experiment 2b, there was no significant change in freezing to S1 across second-order conditioning (Figure 2F, *F*_(1, 23)_ = 2.12, *p* > 0.05).

At test, rats infused with CHX showed significantly less freezing when tested with S1 than rats in group VEH (Figure 2C, *F*_(1, 46)_ = 17.77, *p* < 0.001, Figure 2E, *F*_(1, 19)_ = 7.24, *p* < 0.05, Figure 2G, *F*_(1, 23)_ = 5.30, *p* < 0.05). This was true regardless of whether the CHX infusion occurred before or after first-order conditioning or after second-order conditioning. In all cases, there was no significant effect of CHX on levels of freezing to the S2 (*F*s < 1, *p* > 0.05).

### Experiment 3

Experiment 2 found no evidence to support the hypothesis that second-order conditioned fear of the S2 is consolidated via proteins synthesized during first-order conditioning of the S1. Infusion of the protein synthesis inhibitor cycloheximide into the BLA consistently disrupted consolidation of fear to the S1 while leaving intact fear to the S2. Experiment 3 examined whether this contrasting effect of cycloheximide on consolidation of fear to the S2 and S1 is specific to the truncated second-order protocol. To do so, we reversed the order of exposure to the S1-shock and S2àS1 pairings in stages 1 and 2 to generate a truncated sensory preconditioning protocol. We then used this protocol to examine the effect of a BLA cycloheximide infusion on consolidation of first-order fear to the S1 and sensory preconditioned fear to the S2.

Conditioning to the S1 was successful across stage 2. There was a significant linear increase in freezing to the S1 across S1-shock pairings (Figure 3B, *F*_(1,16)_ = 35.83, *p* < 0.001). There were no between-group differences in the development of freezing across the pairings or in the overall levels of freezing (*F*s < 1).

**Figure 3.**
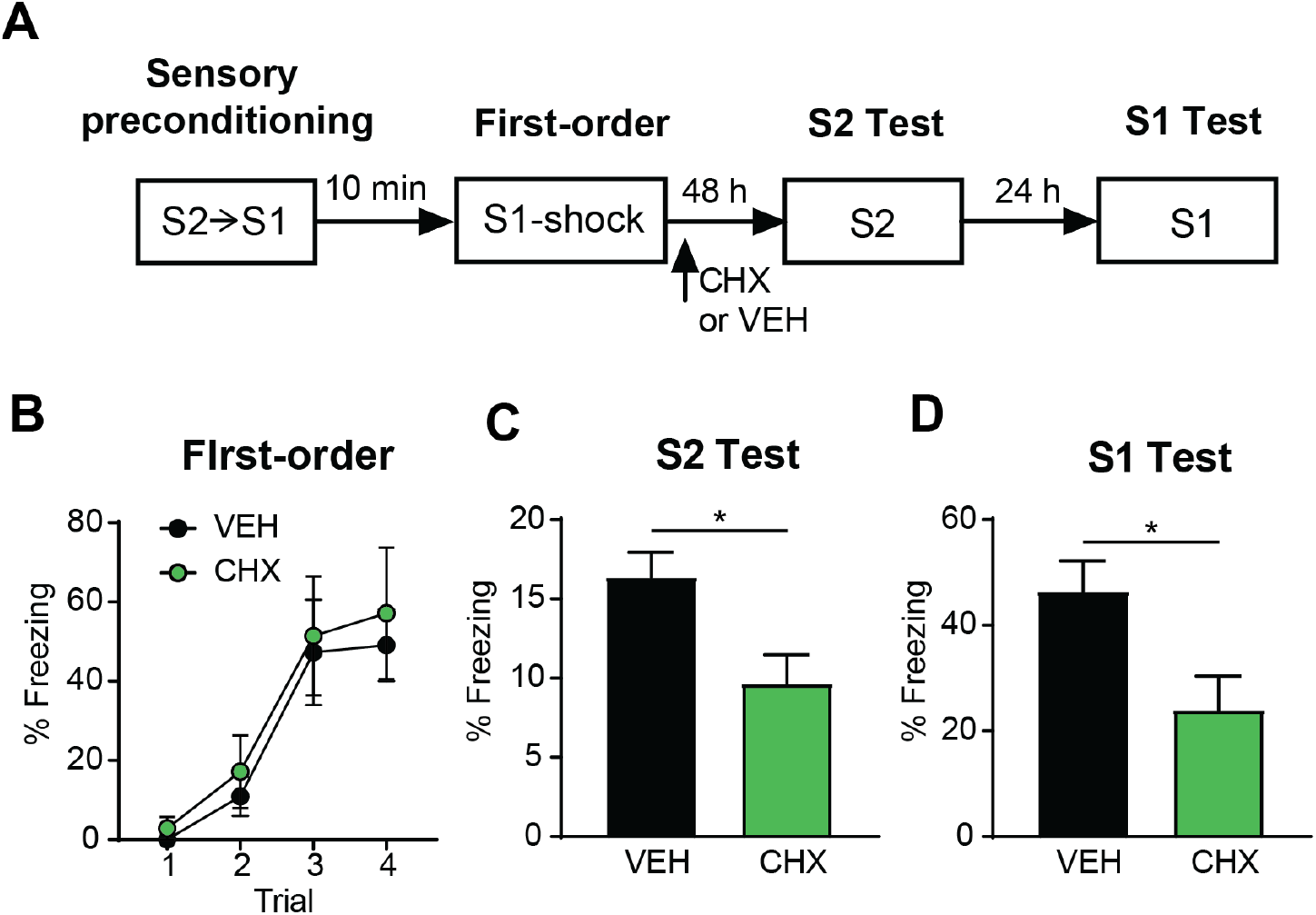
Protein synthesis in the BLA is required for consolidation of fear to S1 and S2 in the truncated sensory-preconditioning protocol. (A) Experimental timeline of conditioning and testing for rats that received intra-BLA infusions of CHX or VEH. (B) Mean (±SEM) levels of freezing to presentations of the S1 during first-order conditioning. (C) Mean (±SEM) levels of freezing to presentations of the S2 and the S1 during second-order conditioning. (D-E) Mean (+SEM) levels of freezing across test presentations of the S2 (D) and the S1 (E).

At test, rats in Group CHX froze significantly less than control rats across test presentations of S1 (Figure 3D, *F*_(1,16)_ = 6.35, *p* < 0.05) and the S2 (Figure 3C *F*_(1,16)_ = 7.42, *p* < 0.05). This contrasts with the previous experiments where cycloheximide failed to disrupt consolidation of the S2. Taken together, these results indicate that absence of a cycloheximide effect in the previous experiments was not simply due to S2 being subjected to higher-order conditioning. Rather, it is a consequence of something specific to second-order conditioning.

### Experiment 4

The next series of experiments examined the possibility that the protein synthesis requirement for consolidating fear to S2 depends on the content of the association that forms during second-order conditioning. Past research has shown that the content of the association formed during second-order conditioning depends on the temporal relation between S2 and S1. When these two stimuli are presented in a serial compound (S2àS1), as in our standard protocol, responding to the S2 is unaffected by extinction of the S1 or habituation to the US (Rizley and Rescorla, 1972; Holmes et al., 2014). This suggests that responding to the S2 is mediated by an S-R association between the S2 and fear responses elicited by the S1. In contrast, when S2 and S1 are presented in simultaneous compound (S2S1), fear to the S2 is eliminated by extinction of the S1 (Rescorla, 1982). That is, responding to S2 is contingent on the current value of the S1, indicating that rats formed an S-S association.

The next experiments used the serial and simultaneous protocols to test the hypothesis that protein synthesis in the BLA is required to consolidate fear mediated by S-S associations but not to consolidate fear mediated by S-R associations. We began by demonstrating that fear acquired to the S2 in the simultaneous protocol is contingent upon its pairing with S1 and the prior pairings of the S1 and shock. In order to test this possibility, we used three groups of rats. Rats in Group PP were exposed to S1-shock pairings in stage 1 and to presentations of the simultaneous S2S1 compound in stage 2. Rats in Group PU were exposed to S1-shock pairings in stage 1 and to explicitly unpaired presentations of S2 and S1 in stage 2. Rats in Group UP were exposed to unpaired presentations of S1 and shock in stage 1 and to presentations of the simultaneous S2S1 compound in stage 2.

Figure 4B shows the levels of freezing to S1 during stage 1. It shows that freezing to S1 increased across trials (*F*_(1, 21)_ = 129.45, *p* < 0.001). The rate of acquisition of freezing to S1 was different among the three groups (linear trial x group interaction: *F*_(1, 21)_ = 23.97, *p* < 0.001): only rats exposed to S1-shock pairings (Groups PP and PU) showed increased freezing to S1 across trials. This was also reflected in the overall levels of freezing, where Groups PP and PU froze significantly more to S1 than rats in Group UP (*F*_(1, 21)_ = 31.68, *p* < 0.001) but did not differ from each other (*F*_(1, 21)_ < 1.3).

**Figure 4.**
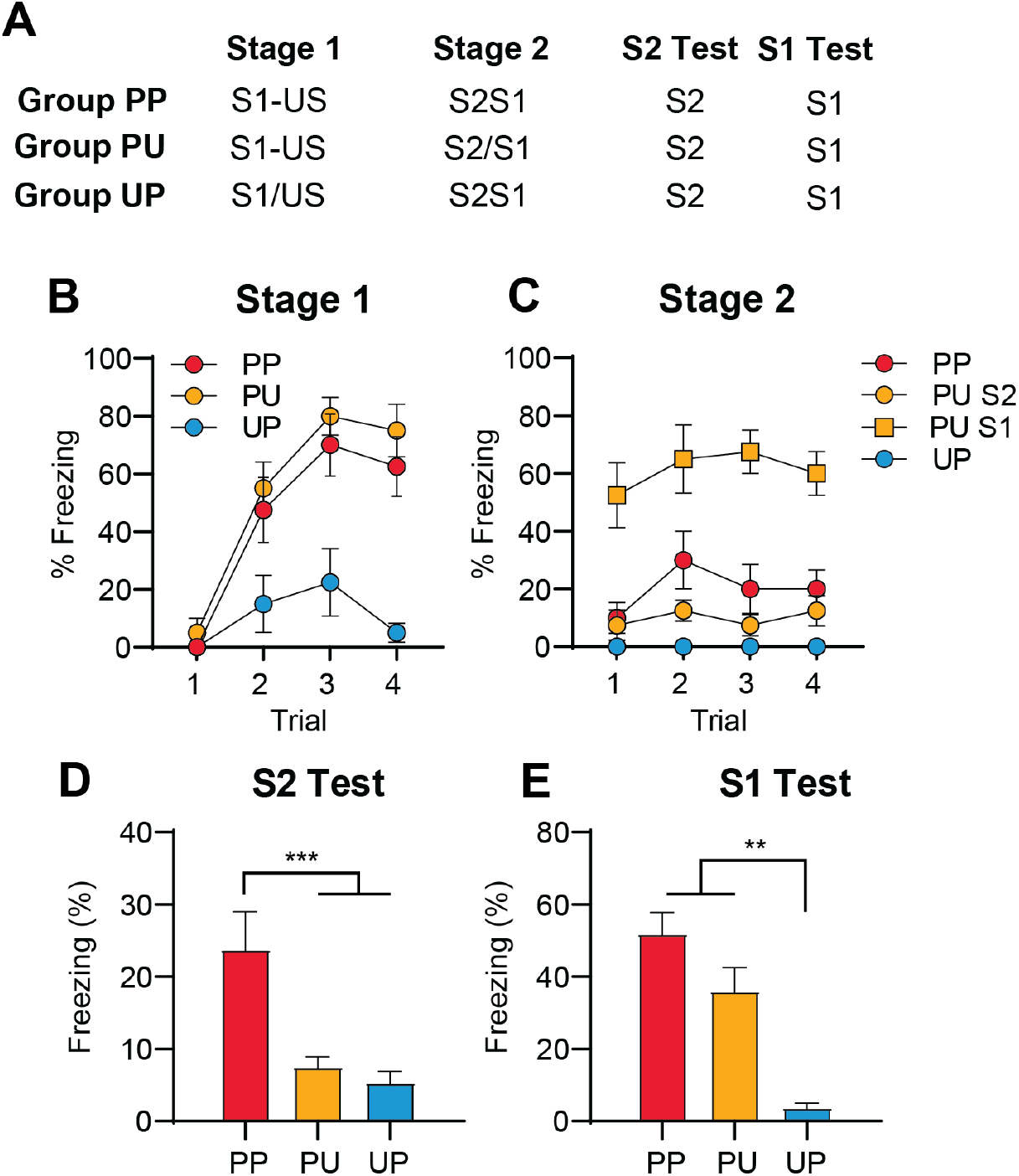
Freezing to S2 in the simultaneous protocol is due to second-order conditioning. (A) Experimental groups for conditioning and testing. A (/) indicates that the stimuli were explicitly unpaired (B) Mean (±SEM) levels of freezing to presentations of the S1 during stage 1. (C) Mean (±SEM) levels of freezing to presentations of the S2 and S1 during stage 2. (D) Mean (+SEM) levels of freezing across test presentations of the S2.

Figure 4C shows the levels of freezing to S2 and S1 during stage 2. S2 and S1 were presented simultaneously for Groups PP and UP, hence for these groups only one dataset is presented for freezing to S2 and S1 in Figure 6C. For Group PU, the S2 and S1 were presented separately in stage 2 and, hence, separate datasets are presented for freezing to S2 and S1. Freezing to the S2 and S1 was analyzed separately, with the data for rats in Groups PP and UP being duplicated across the two analyses. To account for this duplication, the criterion for rejection of the null hypothesis (α) was reduced to 0.025 for analysis of stage 2 training only.

During presentations of the S2, Group PP froze significantly more than rats in Groups PU and UP (*F*_(1, 21)_ = 13.622, *p* = .001), while rats in Groups PU and UP did not differ significantly (*F*_(1, 21)_ = 4.541, *p* = .045). During presentations of the S1, groups that received pairings of S1 and shock (Groups PP and PU) froze significantly more than rats in Group UP (*F*_(1, 21)_ = 47.68, *p* < .001). Further, rats in Group PU froze significantly more to S1 than rats in Group PP, who were exposed to S1 in compound with S2 (*F*_(1, 21)_ = 41.58, *p* < .001). This is likely due to external inhibition of freezing to S1 by the novel S2, or a generalization decrement from the conditioned S1 to its presentation in compound with S2.

When rats were tested for second-order conditioned fear (Figure 6D), Group PP froze significantly more than the other groups (*F*_(1, 21)_ = 18.790, p < 0.001), while Groups PU and UP did not differ significantly (*F*_(1, 21)_ < 1.8). Figure 6E shows test levels of freezing to S1. Rats that had been exposed to S1-shock pairings in stage 1 (Groups PP and PU) froze significantly more than rats that had been exposed to unpaired presentations of the S1 and shock (Group UP; *F*_(1, 21)_ = 15.93, *p* = 0.01); and Groups PP and PU did not differ significantly (*F*_(1, 21)_ = 0.224, *p* = 0.641). These results indicate that freezing to S2 in the simultaneous protocol is due to second-order conditioning rather than generalization of fear from S1, or any unconditioned ability of S1 to condition freezing to S2.

### Experiments 5a-b

Experiments 5a and 5b sought to replicate previous demonstrations (Rescorla, 1982, Holmes et al., 2014) that extinction of the previously conditioned S1 differentially affects responding to S2 at test depending on whether S2 and S1 had been presented in a serial (S2àS1) or simultaneous (S2S1) compound during second-order conditioning. Rats received S1-shock pairings in stage 1 and pairings of S2 and S1 in stage 2. The experiments differed with respect to the presentation of the stimuli during second-order conditioning in stage 2. In Experiment 5a, rats received serial S2àS1 pairings, whereas in Experiment 5b, rats received simultaneous presentations of S2 and S1. The following day, rats in group EXT received twenty S1 alone exposures, while rats in group No EXT received an equivalent amount of context exposure. All groups were then tested for fear to the S2 and the S1.

The results of Experiments 5a and 5b are presented in Figure 5. All rats acquired first-order fear to the S1 in stage 1 and second-order fear to the S2 in stage 2. Freezing to the S1 increased across the four S1-shock pairings (Figure 5B, *F*_(1, 14)_ = 376.45, *p* < 0.001, Figure 5G, (*F*_(1, 16)_ = 871.76, *p* < 0.001) and there were no significant between-group differences or group x trend interactions (*Fs* < 2, *p* > 0.05). During stage 2, rats in the serial second-order groups showed an increase in freezing to the S2 across the S2àS1 pairings (Figure 5C, *F*_(1, 14)_ = 46.70, *p* < 0.001), while freezing to the S1 remained stable (*Fs* < 1.44, *p* > 0.05). Rats in the simultaneous groups showed a significant increase in freezing across simultaneous presentations of S2 and S1 (Figure 5H, *F*_(1, 16)_ = 6.159, *p* <). In both experiments, there were no significant between-group differences or group x trend interactions during stage 2 (*Fs* < 1.20, *p* > 0.05).

**Figure 5.**
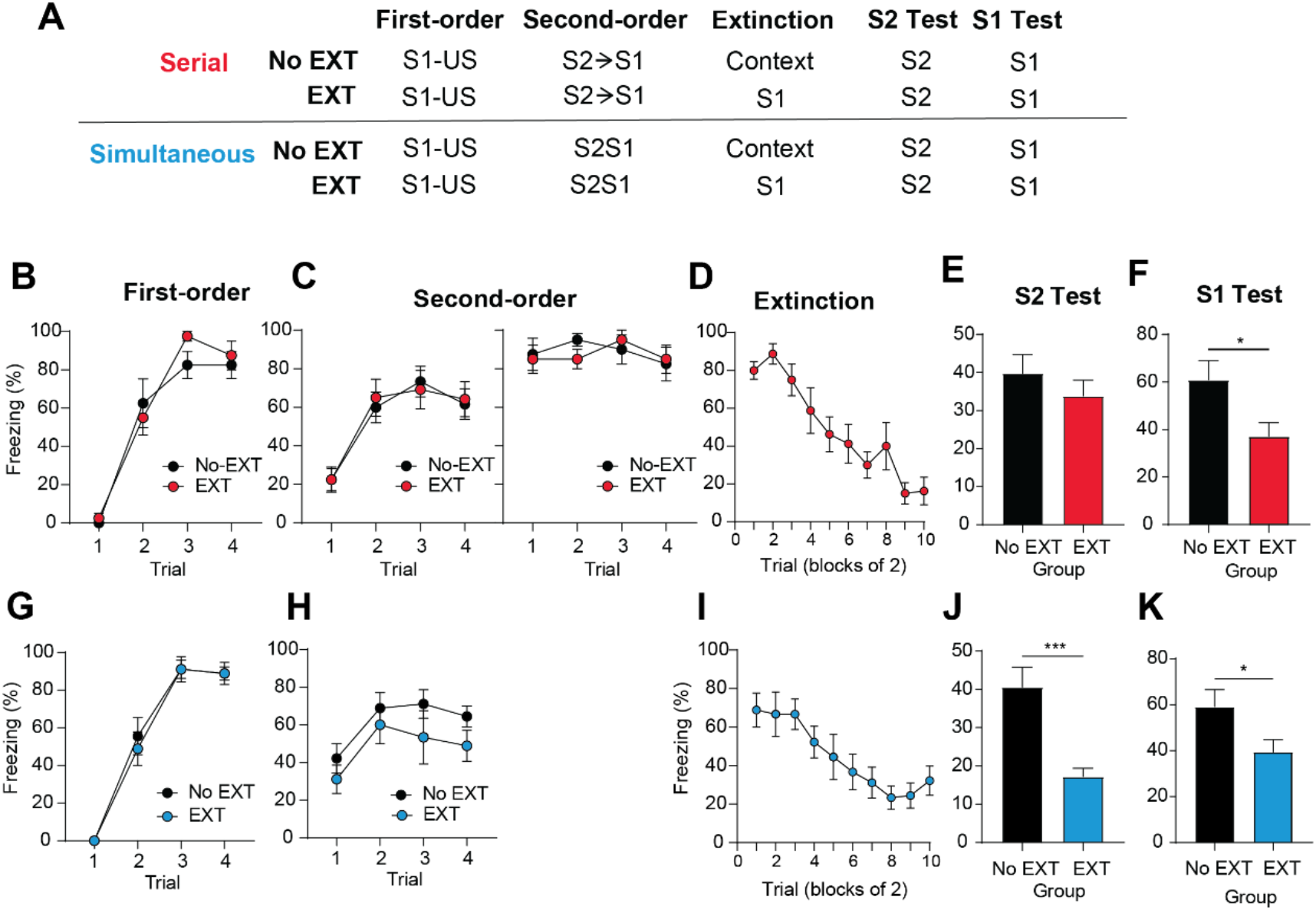
Extinction of S1 reduces responding to S2 in the simultaneous second-order conditioning protocol (S2S1), but not in the serial second-order conditioning protocol (S2àS1). (A) Experimental groups for conditioning and testing. (B) Mean (±SEM) levels of freezing to presentations of the S1 during first-order conditioning. (C) Mean (±SEM) levels of freezing to presentations of the S2 and the S1 during second-order conditioning. (D) Mean (+SEM) levels of freezing across test presentations of the S2.

**Figure 6.**
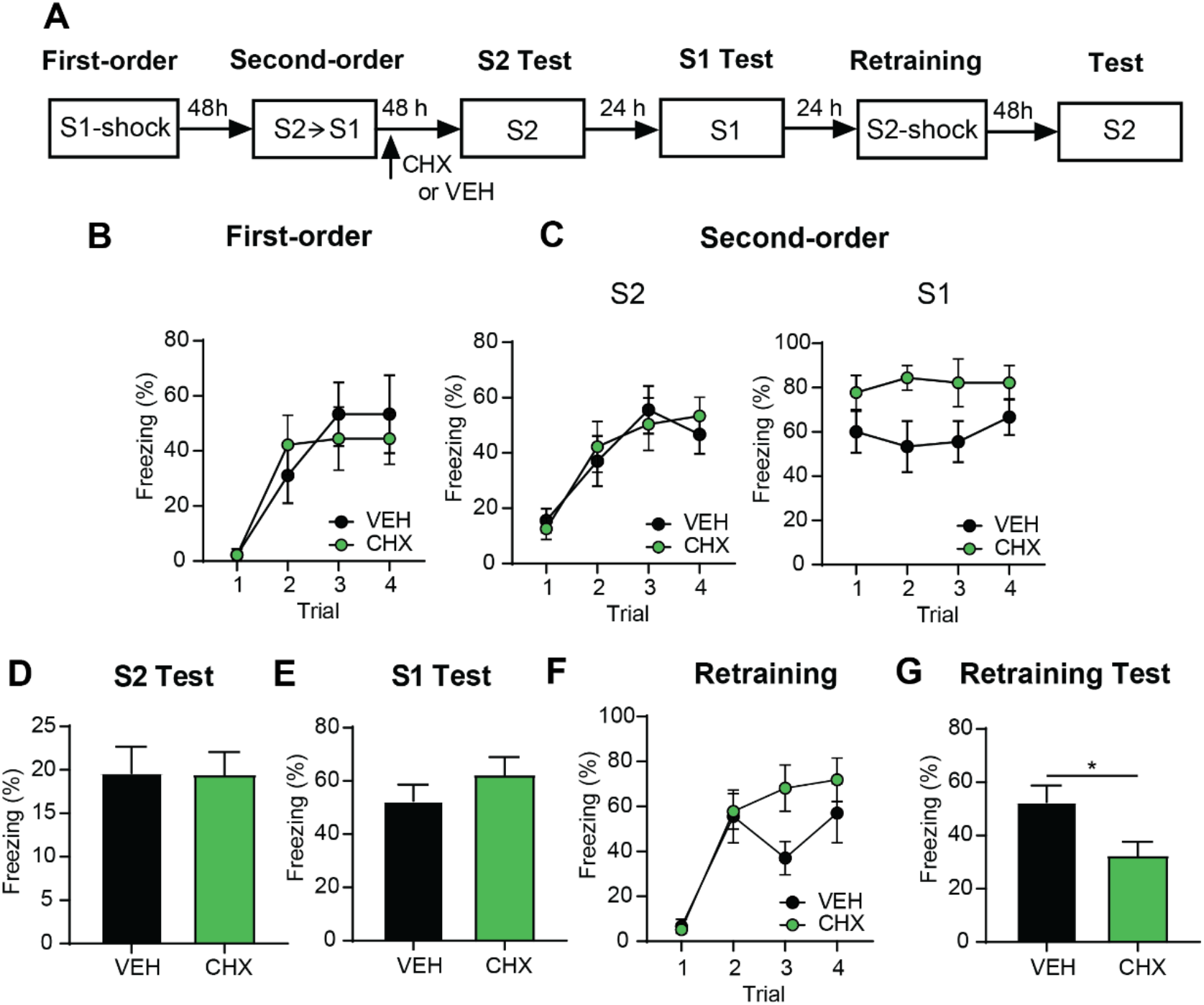
Consolidation of fear to S2 in the serial second-order conditioning protocol does not require *de novo* protein synthesis in the BLA. (A) Experimental timeline of conditioning and testing for rats that received intra-BLA infusions of CHX or VEH. (B) Mean (±SEM) levels of freezing to presentations of the S1 during first-order conditioning. (C) Mean (±SEM) levels of freezing to presentations of the S2 and the S1 during second-order conditioning. (D-E) Mean (+SEM) levels of freezing across test presentations of the S2 (D) and S1 (E). (F-G) Mean (±SEM) levels of freezing to presentations of the S2 during its reconditioning (F) and testing (G).

During extinction training, there was a significant decrease in freezing across presentations of the S1 (Figure 5D, *F*_(1, 7)_ = 91.56, *p* < 0.001, Figure 5I, *F*_(1, 8)_ = 23.132, *p* < 0.001). At test, rats in Group Simultaneous EXT showed significantly lower freezing to both the S2 and S1 than rats in Group Simultaneous No EXT (*F*_(1, 16)_ = 16.84, *p* < 0.001). In contrast, levels of freezing to the S2 did not differ between Groups Serial EXT and Serial No EXT (*F*_(1, 14)_ = 0.85, *p* = 0.37). This was not due to a failure to extinguish the S1, as test levels of freezing to the S1 were significantly lower in Group Serial EXT compared to Group Serial No EXT (*F*_(1, 14)_ = 5.8, *p* < 0.05). These results replicate previous findings demonstrating that conditioned responding to the S2 is sensitive to extinction of its first-order associate, S1, when S2 and S1 are presented in simultaneous compound during second-order conditioning, but not when they are presented serially.

### Experiment 6

Experiments 6a and 6b examined whether protein synthesis in the BLA is needed to consolidate fear to the S2 in the serial and simultaneous second-order conditioning protocols. In both experiments, rats were exposed to S1-shock pairings in stage 1. In Experiment 6a, S2 was conditioned in serial-order with the previously conditioned S1 (S2àS1). In Experiment 6b, S2 and S1 were presented simultaneously during second-order conditioning (S2S1). In both experiments, rats received an intra-BLA infusion of cycloheximide or vehicle immediately after stage 2. Rats were subsequently tested for fear to both the S2 and S1.

### Experiment 6a

Figure 6 shows levels of freezing during first-order conditioning. Averaged across groups, there was a significant increase in freezing across S1-shock pairings (*F*_(1, 16)_ = 21.29, p < 0.001). There was no significant difference in freezing levels between rats assigned to the CHX and VEH groups (*F*_(1, 16)_ = 0.031, p = 0.862) and there was no significant group x trend interaction (*F*_(1, 16)_ = 0.5, *p* = 0.49).

In stage 2, freezing levels to the S2 increased across conditioning trials (*F*_(1, 16)_ = 52.00, p < 0.001). Freezing levels to the S2 did not differ between CHX and VEH groups (*F*_(1, 16)_ = 0.012, p= 0.914) and there was no significant group x trend interaction (*F*_(1, 16)_ = 0.304, p= 0.589). Freezing levels to the S1 during second-order conditioning were significantly higher in the rats assigned to the CHX group compared to the VEH group (*F*_(1, 16)_ = 5.462, *p* < 0.05). There was no significant effect of trial (*F*_(1, 16)_ = 0.655, p = 0.430) or group x trial interaction (*F*_(1, 16)_ = 0.073, *p* = 0.790) on levels of freezing to the S1.

Consistent with previous research (Lay et al., 2018; Leidl et al., 2018), both groups froze at similar levels to both the S2 and the S1 at test. Statistical analyses showed that there was no significant effect of CHX on levels of freezing to either the S2 (*F*_(1, 16)_ = 0.002, p = 0.965) or the S1 (*F*_(1, 16)_ = 1.136, *p* = 0.302).

In order to confirm that the BLA infusion of CHX was effective in our experiment, we sought to replicate, within the same animals, the well-documented effect of CHX on consolidation of first-order conditioned fear. Thus, S2 was completely extinguished after initial S1 and S2 testing and then retrained as a first-order stimulus by exposing rats to four S2-shock pairings. Immediately after first-order retraining, rats were infused with CHX or VEH and then tested for fear to the S2 the following day. Figure 6F-G shows levels of freezing to the S2 across its pairings with shock and subsequent testing of the S2. There was a significant increase in freezing to the S2 across its pairings with shock (*F*_(1, 16)_ = 46.531, *p* < 0.001) and no significant effect of group (*F*_(1, 16)_ = 1.55, *p* = 0.231) or group x trend interaction (*F*_(1, 16)_ = 2.39, *p* = 0.141). At test, there was significantly higher freezing in the VEH group compared to the CHX group (*F*_(1, 16)_ = 6.140, *p* < 0.05), indicating the drug had successfully disrupted consolidation of first-order conditioned fear to the S2.

### Experiment 6b

Figure 7 shows levels of freezing during first-order conditioning. Freezing to the S1 increased across S1-shock pairings (*F*_(1, 20)_ = 201.31, *p* < 0.001) and this increase did not differ across groups (*F*_(1, 20)_ = 0.040, *p* = 0.84). Overall levels of freezing to the S1 did not differ between rats allocated to the CHX and VEH groups (*F*_(1, 20)_ = 0.03, *p* = 0.86). During second-order conditioning, freezing to the S2S1 simultaneous compound did not differ across pairings or groups and there was no significant group x trial interaction (*Fs* < 1.96, *p* > 0.05).

**Figure 7.**
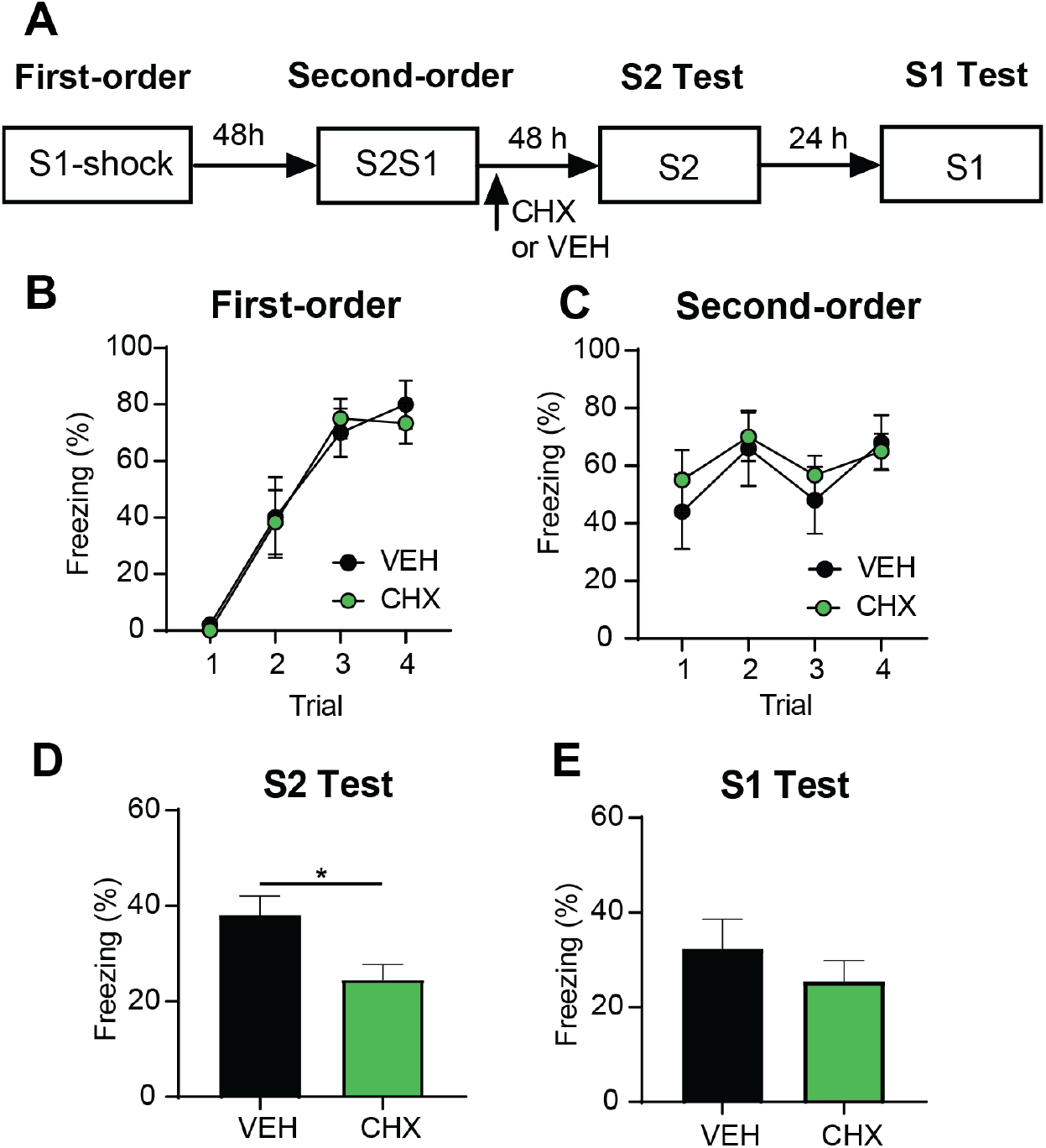
Consolidation of fear to S2 in the simultaneous second-order conditioning protocol requires *de novo* protein synthesis in the BLA. (A) Experimental timeline of conditioning and testing for rats that received intra-BLA infusions of CHX or VEH. (B) Mean (±SEM) levels of freezing to presentations of the S1 during first-order conditioning. (C) Mean (±SEM) levels of freezing to simultaneous presentations of S2 and S1 during second-order conditioning. (D-E) Mean (+SEM) levels of freezing across test presentations of the S2 (D) and the S1 (E).

**Figure 8.**
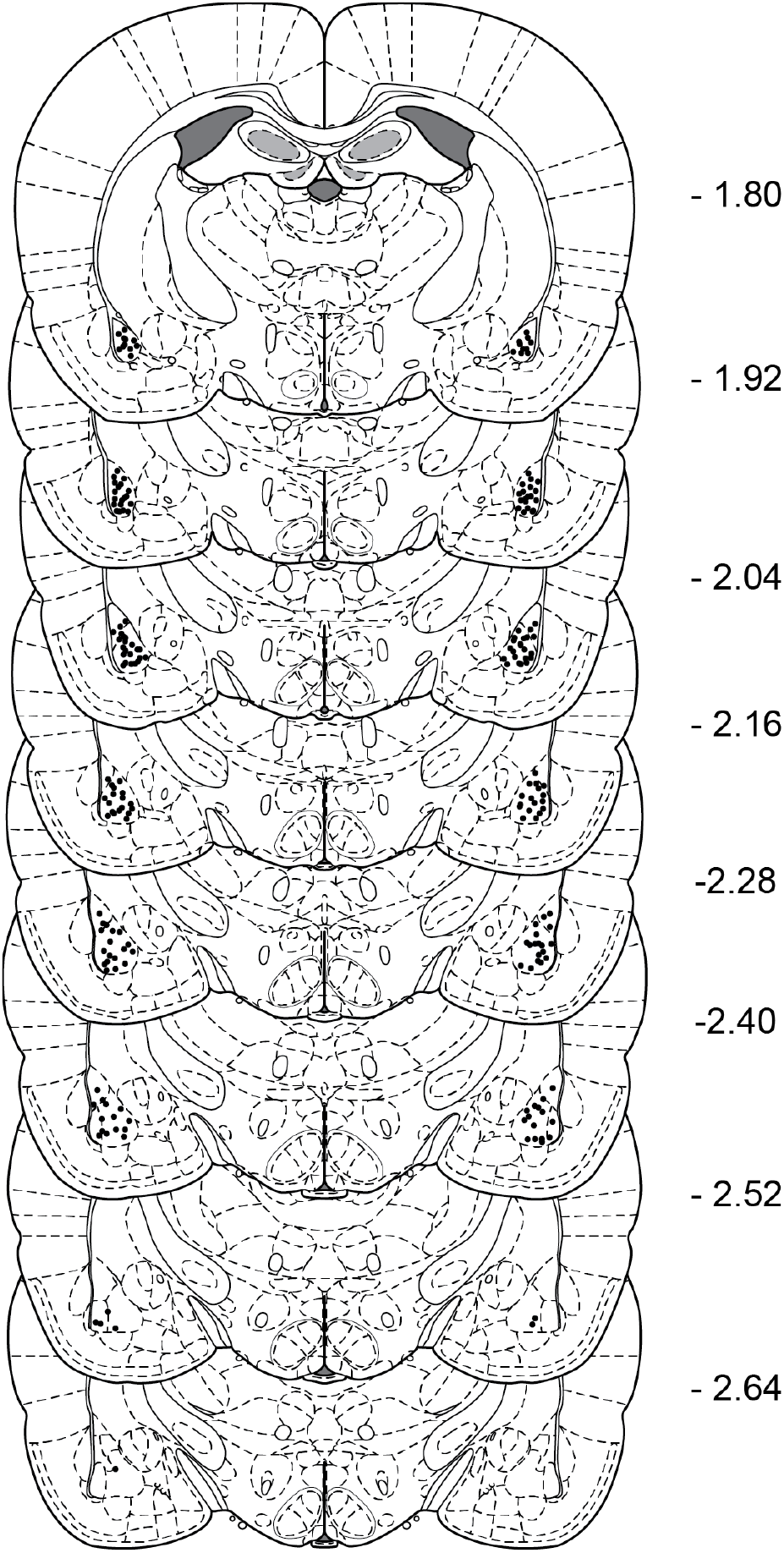
Cannula placements for all rats included in the study. The black dot represents the most ventral position of the cannula.

At S2 test, there was significantly lower freezing in Group CHX compared to Group VEH (*F*_(1, 20)_ = 6.99, *p* < 0.05). Freezing to the S1 did not differ significantly between groups VEH (*F*_(1, 20)_ = 0.82, *p* = 0.38). Thus, consolidation of fear to S2 in the simultaneous second-order protocol *does* require protein synthesis in the BLA.

## Discussion

The present study originated in recent findings that protein synthesis in the BLA is required to consolidate first- but not second-order conditioned fear (Lay et al., 2018; Leidl et al., 2018). The aim was to determine why this is the case and, thereby, the circumstances under which protein synthesis is needed to consolidate new information in the BLA. In each experiment, rats were exposed to a session of S1-shock pairings in stage 1 and a session of S2-S1 pairings (serial, S2àS1; or simultaneous, S2S1) in stage 2. The latter session was preceded or followed by an intra-BLA infusion of the protein synthesis inhibitor, cycloheximide. Finally, rats were tested for fear (measured in freezing) to test presentations of the S2 alone and S1 alone.

The first set of experiments tested the hypothesis that consolidation of second-order fear can be supported by proteins that were translated during first-order conditioning: hence, it does not require *de novo* protein synthesis in the BLA. One way this could occur is via a mechanism of the sort described in “tag-and-capture” theories (Frey and Morris, 1997; Frey and Morris, 1998a; Frey and Morris, 1998b; Redondo and Morris, 2011). According to these theories, the synapses that respond to an event (e.g., CS-US pairings) are initially tagged in some way; and this tag permits capture of plasticity-related proteins for stabilization of the synaptic response. Applied to second-order conditioning, the idea is that S2 could capture proteins synthesized during the prior conditioning of S1, resulting in protein synthesis-independent strengthening of synapses that respond to the S2.

To test this hypothesis, we examined the effect of a BLA cycloheximide infusion on consolidation of second-order fear using a protocol in which the interval between S1-shock pairings in stage 1 and S2àS1 pairings in stage 2 was just 10 minutes (as opposed to 48 h used previously). Under these circumstances, consolidation of second-order fear could not be supported by proteins synthesised during first-order conditioning as these would also be disrupted by the cycloheximide infusion: thus, if the hypothesis is correct, cycloheximide should disrupt fear to both the S1 and S2 at test. The initial experiment confirmed that freezing to S2 in this “truncated” protocol is indeed due to second-order conditioning (Exp 1): rats exposed to paired stimulus presentations in both stages of training froze more when tested with S2 than rats exposed to unpaired stimulus presentations in either stage of training. Subsequent experiments then showed that, contrary to the hypothesis, the effects of cycloheximide on the test levels of freezing to S1 and S2 were completely dissociable: the drug disrupted freezing to S1 without affecting freezing to S2; and this was true regardless of whether it was infused into the BLA before the session of S1-shock pairings in stage 1 (Exp 2c), or before/after the session of S2àS1 pairings in stage 2 (Exp 2a, 2b). Together, these findings confirm that consolidation of first-order fear is a time- and protein synthesis-dependent process in the BLA (note the absence of any acute drug effects on freezing to S1 in Fig. 2D [Exp 2b] and Fig. 2F [Exp 2c]); and show that consolidation of second-order fear does *not* depend on proteins synthesized during first-order conditioning. That is, second-order fear to S2 was intact even through first-order fear to S1 was disrupted, showing that consolidation of fear to S2 simply does not require protein synthesis in the BLA.

The second set of experiments examined whether the differential protein synthesis requirement for consolidation of first- and second-order fear is related to differences in what is learned in each of these protocols. Specifically, we tested the hypothesis that protein synthesis in the BLA is needed to consolidate S-S associations of the sort that form in first-order conditioning but is *not* needed to consolidate S-R associations of the sort that form in second-order conditioning, at least as it is standardly conducted. To do so, we used two second-order conditioning protocols that have been shown to produce different associations: the standard protocol in which S2 and S1 are presented serially in stage 2 (S2àS1); and another protocol in which S2 and S1 are presented simultaneously in stage 2 (S2S1). The initial experiments confirmed that the two protocols generate different associations. Fear to S2 in the simultaneous protocol was reduced by extinction of the S1 indicating that, in this case, S2 associates with S1 across the S2S1 pairings (Exp 4, 5a) and comes to elicit fear through integration of S2-S1 and S1-shock associations at test. By contrast, fear to S2 in the serial protocol was unaffected by extinction of the S1. This is consistent with the well-established view that, in this case, S2 associates with the fear elicited by S1 across the S2àS1 pairings (Exp 5b) and, thereby, elicits fear directly at test (via an S2-fear [S-R] association) (Rizley & Rescorla, 1972, Rescorla, 1973b).

Subsequent experiments then examined the protein synthesis requirement for consolidation of fear to S2 in the simultaneous and serial protocols. They showed that a BLA infusion of cycloheximide disrupted consolidation of fear to S2 in the simultaneous protocol: relative to controls, rats that received this infusion after the session of S2S1 pairings froze less when tested with S2 but just as much when tested with the S1, suggesting a specific disruption of the S2-S1 association (Exp 6). By contrast, a BLA cycloheximide infusion had no effect on consolidation of the S2-fear association in the serial protocol: rats that received this infusion after the session of S2àS1 pairings froze just as much as the controls when tested with both S2 and S1 (Exp 7). Thus, within the BLA, the protein synthesis requirement for consolidation of Pavlovian conditioned fear does indeed depend on what is learned in conditioning. Protein synthesis is needed to consolidate fear to S2 when it is supported by integration of S2-S1 and S1-shock associations in the simultaneous protocol; but is not needed to consolidate fear to S2 when it is supported by an S2-fear association in the serial protocol.

Why is protein synthesis in the BLA needed to consolidate fear to S2 in the simultaneous protocol but not in the serial protocol? While the BLA is the site where protein synthesis is needed to integrate/consolidate the S2-S1 and S1-shock associations in the simultaneous protocol, the protein synthesis requirement for consolidation of the S2-fear association in the serial protocol may lie elsewhere in the circuitry engaged for fear expression. Obvious candidate regions include the central nucleus of the amygdala (CeA), periaqueductal grey (PAG) and prelimbic region of the medial prefrontal cortex (PL). Each of these regions is strongly connected with the BLA (which plays a role in consolidating the S2-fear association) and has been implicated in maintaining/expressing different components of a fear response (autonomic, behavioral, endocrine) (Di Scala et al., 1987; Walker and Carrive, 2003; Corcoran and Quirk, 2007; Ciocchi et al., 2010; Tovote et al., 2015). However, at present, only the CeA has been examined for its involvement in second-order conditioned fear (see Holmes et al., 2022). Lay et al (2018) showed that protein synthesis in the CeA is not required to consolidate fear of S2 in the serial protocol used here: rats that received a CeA infusion of cycloheximide immediately after the session of S2àS1 pairings exhibited just as much freezing to S2 at test as the vehicle-infused controls. As such, future work will examine the involvement of the PAG and/or PL in consolidating second-order fear to S2: specifically, whether this consolidation requires protein synthesis-dependent changes in the PAG and/or PL; and, if so, how these changes might be regulated by the BLA. We hypothesize that the BLA coordinates consolidation of the S2-fear association by regulating protein synthesis-dependent changes in the PAG and/or PL; and will seek to determine how this is achieved via the known requirements for CaMK signaling, gene transcription and DNA methylation in the BLA (Lay et al., 2018).

In summary, the present study has shown that, within the BLA, the protein synthesis requirement for consolidation of second-order fear is determined by what is learned when the S2 is paired with the already-conditioned S1. Protein synthesis is required to consolidate fear to S2 when it is supported by integration of S2-S1 and S1-shock associations formed in training; but is not required to consolidate fear to S2 when it is supported by a direct association between S2 and the central state of fear elicited by the S1 (S2-fear). These findings are significant as they show that the substrates of learning and memory in the mammalian brain are not fixed and immutable; instead, these substrates vary with the information that animals extract from their environments. Future research will examine how direct stimulus-fear associations are consolidated in the brain: specifically, whether circuitry that underlies expression of fear responses is involved in acquisition of stimulus-fear associations; and how the BLA interacts with this circuitry to consolidate the stimulus-fear associations.

## Methods

### Surgery

All rats were surgically implanted with bilateral cannulas targeting the BLA. Rats were anesthetized with isoflurane gas (5% for induction, 2-2.5% for maintenance; Cenvet) and mounted on a stereotaxic apparatus. Two 26-gauge guide cannulas were implanted into the brain of each rat through two holes drilled in the skull. The tips of the guide cannulas were aimed at the BLA in each hemisphere (2.4 mm posterior to Bregma, 4.9 mm lateral to the midline and 8.3 mm ventral to Bregma). The guide cannulas were secured in position with dental cement and four jeweler screws. A dummy cannula was kept in each guide at all times other than during infusions. Immediately after surgery, rats received an intraperitoneal (i.p.) injection of penicillin (0.3 mL) and were placed on a heating mat until they had recovered from the effects of the anesthetic. They were then returned to their home cages. Rats were given a minimum of seven days to recover from of surgery. During this period, rats were handled and weighed daily.

### Drug infusions

Cycloheximide is commonly used to block protein synthesis in discrete regions of the brain (Schneider-Poetsch et al., 2010), and its infusion into the BLA disrupts consolidation of first-order conditioned fear (e.g., Duvarci, 2005). Cycloheximide (Sigma-Aldrich) was dissolved in 70% ethanol to yield a stock solution with 200 ug/uL concentration. This was then diluted 1:4 with artificial cerebral spinal fluid (ACSF; Sigma Aldrich) to a final concentration of 40 ug/uL. Vehicle was made by diluting 70% ethanol 1:4 with ACSF (Duvarci et al., 2005).

Rats received bilateral BLA infusions of cycloheximide (CHX) or vehicle (VEH; 0.5 uL per hemisphere). The infusion procedure for each rat commenced with removal of dummy caps from the two guide cannulas and insertion of 33-gauge internal cannulas. The two internal cannulas were each connected to separate 25 uL Hamilton syringes, which were driven by an infusion pump (Harvard Apparatus). Drug or vehicle was infused bilaterally into the BLA through the internal cannulas at a rate of 0.25 uL/min. Following infusion, the infusion cannulas were left in place for an additional two minutes to allow for diffusion of the drug away from the cannula tip.

### Histology

Following behavioral testing, rats were euthanized with a lethal dose of sodium pentobarbital (injected i.p.). Rats were decapitated, their brains rapidly removed and frozen. Brains were sliced on a cryostat into coronal sections of 40 um thickness. Every other section through the region of interest was mounted on a glass microscope slide and then stained with cresyl violet. Slides were observed under a microscope to confirm the location of cannulas using the brain atlas of Paxinos and Watson (2007). Rats with inaccurate cannula placements (one or both cannulas positioned outside the boundaries of the BLA) or extensive damage to the BLA were excluded from the statistical analysis.

### Behavioral procedures

#### Experiment 1

##### Contextpre-exposure

All rats were familiarized with the context on days 1 and 2. There were two sessions of context pre-exposure on each of those days, spaced three hours apart, with each session lasting twenty minutes. This was done to eliminate any neophobic context responses that could interfere with detection of conditioned responses.

##### First-order conditioning

On day 3, rats in Groups PP and PU received four pairings of S1 and a 0.8 mA x 0.5 s foot shock. The duration of S1 was 10 s and its presentation coterminated with the foot shock. Onset of the first S1-shock pairing occurred five min after placement in the chamber, and each subsequent pairing five min later (measured from the offset of the previous stimulus presentation). Rats remained in the chamber for an additional one min after the final S1-shock pairing. Rats in Group UP received equivalent exposure to S1 and shock but presented in an explicitly unpaired arrangement. S1 and shock were presented in the order S1, shock, shock, S1, S1, shock, shock, S1, with an ITI of three min and the first S1 presentation occurring two min after placement in the chamber. Rats remained in the chamber for one min following the final S1 presentation.

##### Second-order conditioning

Rats were returned to the chambers 10 min later for stage 2. Rats in Groups PP and UP received four pairings of a 30 s S2 with the already conditioned S1. Here, after a two min baseline period, S2 was presented and followed immediately by S1. This pairing was repeated three times with a five min ITI, yielding a total of four S2àS1 pairings. Rats in Group PU received four presentations each of the S2 and S1 (i.e., eight stimulus presentations in total). The first stimulus presentation occurred two min after placement in the chamber and the interval between stimulus presentations was fixed at 2.5 min. S2 and S1 were presented in the order S2, S1, S1, S2, S2, S1, S1, S2.

##### Context extinction

Rats received context extinction on day 4 in order to reduce the level of freezing in the context alone and, thereby, ensure that the subsequent measure of freezing to S2 was not confounded with freezing elicited by the context. Rats were placed in the chambers for two sessions in the absence of any stimuli. Each session was 20 min in duration and spaced approximately three hours apart. Rats received a further 10 min extinction exposure to the chambers on the morning of Day 5. This was done to extinguish any context-elicited freezing that had spontaneously recovered from the previous day and that would have interfered with the detection of freezing to S2.

##### Test

Rats were tested with S2 on the afternoon of day 5 and with S1 on the afternoon of day 6. Rats were placed into the chambers and, following a two min adaptation period, the S2 or S1 was presented. There were eight stimulus presentations in each test session, and the interval between the presentations was fixed at three min. Across the test sessions, the durations of S2 and S1 remained 30 s and 10 s, respectively.

#### Experiments 2a-c

Training at testing procedures were identical to those described for Group PP in Experiment 1. Briefly, rats were pre-exposed to the context on days 1 and 2. On day 3 they were exposed to S1-shock pairings, removed from the chamber for approximately 10 min, returned to the chambers and exposed to pairings of S2 and S1 (S2àS1). On day 4 rats received context extinction and were then tested for freezing to S2 and S1 across days 5 and 6. The order of testing was counterbalanced such that half the rats in each group were tested with S1 on day 5 and with S2 on day 6, while the remainder were tested with S2 on day 5 and with S1 on day 6.

Training and testing procedures were identical for Experiments 2a-c. The experiments differed with respect to the timing of the intra-BLA CHX or VEH infusion. Rats received intra-BLA infusions either after second-order conditioning (Experiment 2a), after first-order conditioning (Experiment 2b) or prior to first-order conditioning (Experiment 2c).

#### Experiment 3

Behavioral procedures were identical to those used in Experiments 2a-c, except that stages 1 and 2 were reversed. In brief, rats were exposed to the chambers on days 1 and 2. They then received four pairings of S2 and S1 on day 3 (S2àS1). Each 30 s presentation of S2 co-terminated in the onset of a 10 s S1. The first pairing occurred five min after placement in the chambers, the interval between pairings was five min and rats were removed from the chambers two min after the final pairing. The chambers were cleaned and approximately 10 min later, rats were returned to the chambers where they received four S1-shock pairings in the manner described previously. The rats were removed and administered a BLA infusion of either cycloheximide (Group CHX) or vehicle (Group VEH). Following a session of context extinction on day 4, rats were tested with S2 or S1 on days 5 and 6 in the manner described previously. The order of testing was counterbalanced, such that half of the rats in each group were tested with S2 on day 5 and S1 on day 6, and the remainder tested with S2 and S1 in the reversed order. Freezing was not scored across pairings of the affectively neutral S2 and S1 during stage 1 (sensory preconditioning), as rats do not freeze to these stimuli.

#### Experiment 4

In the remaining experiments, the interval between the end of training in stage 1 and start of training in stage 2 was 48 h rather than 10 min. In Experiment 4, rats were familiarized to the contexts on days 1 and 2. On day 3, rats in Groups PP and PU received four pairings of S1 and footshock. Rats in Group UP received equivalent exposure to S1 and shock but presented in an explicitly unpaired arrangement. On day 4, rats received two sessions of context extinction, spaced three hours apart, each lasting 20 min. On day 5, rats received second-order conditioning. Rats in Groups PP and UP received four presentations of a 10 s simultaneous S2S1 compound. Rats in Group PU received four presentations each of the S2 and S1 in an explicitly unpaired arrangement. Rats were tested for fear to the S2 and the S1 on days 6 and 7.

#### Experiments 5a-b

Rats were familiarized to the contexts on days 1 and 2, given first-order conditioning on day 3 and context extinction on day 4. On day 5, rats received second-order conditioning. In Experiment 5a, rats received four serial-order pairing of S2 and S1 (S2àS1). The S2 was 30 s and co-terminated in the 10 s S1. In experiment 5b, rats received four pairings of a simultaneous 10s S2-S1 compound. On day 6, rats in Group EXT were exposed to 20 S1 alone presentations. Rats in Group No EXT were placed in the chambers for an equivalent period of time, but no stimuli were presented. Rats were tested for fear to the S2 and S1 on days 6 and 7, respectively.

#### Experiment 6a-b

Rats were trained in the manner described for the No EXT groups in the previous experiment. Briefly, they were exposed to context alone exposure on days 1 and 2, four S1-shock pairings on day 3, two sessions of context extinction on day 4 and S2 and S1 pairings on day 5. Rats in Experiment 6a received four serial-order S2àS1 pairings. Rats in Experient 6b received four presentations of a simultaneous S2S1 compound. Immediately after the second-order conditioning session, rats received a BLA infusion of cycloheximide (Group CHX) or vehicle (Group VEH). Finally, all rats were tested with S2 alone on day 6 and S1 alone on day 7.

## Acknowledgements

This work was supported by an Australian Research Council (ARC) Discovery Grant to NMH and RFW (DP200102969) and an ARC Future Fellowship to NMH (FT190100697).

## Notes

### Competing Interest Statement

The authors have declared no competing interest.

## References

Barnet RC, Arnold HM, Miller RR (1991) Simultaneous conditioning demonstrated in second-order conditioning: Evidence for similar associative structure in forward and simultaneous conditioning. Learning and Motivation 22:253–268.

Ciocchi S, Herry C, Grenier F, Wolff SB, Letzkus JJ, Vlachos I, Ehrlich I, Sprengel R, Deisseroth K, Stadler MB (2010) Encoding of conditioned fear in central amygdala inhibitory circuits. Nature 468:277–282.

Corcoran KA, Quirk GJ (2007) Activity in prelimbic cortex is necessary for the expression of learned, but not innate, fears. Journal of Neuroscience 27:840–844.

Desgranges B, Lévy F, Ferreira G (2008) Anisomycin infusion in amygdala impairs consolidation of odor aversion memory. Brain research 1236:166–175.

Di Scala G, Mana MJ, Jacobs W, Phillips A (1987) Evidence of Pavlovian conditioned fear following electrical stimulation of the periaqueductal grey in the rat. Physiology & behavior 40:55–63.

Duvarci S, Nader K, LeDoux JE (2005) Activation of extracellular signal-regulated kinase–mitogen-activated protein kinase cascade in the amygdala is required for memory reconsolidation of auditory fear conditioning. European Journal of Neuroscience 21:283–289.

Frey U, Morris RG (1997) Synaptic tagging and long-term potentiation. Nature 385:533–536.

Frey U, Morris R (1998a) Weak before strong: dissociating synaptic tagging and plasticity-factor accounts of late-LTP. Neuropharmacology 37:545–552.

Frey U, Morris RG (1998b) Synaptic tagging: implications for late maintenance of hippocampal long-term potentiation. Trends in neurosciences 21:181–188.

Gewirtz JC, Davis M (1997) Second-order fear conditioning prevented by blocking NMDA receptors in amygdala. Nature 388:471–474.

Gewirtz JC, Davis M. (2000) Using Pavlovian Higher-Order Conditioning Paradigms to Investigate the Neural Substrates of Emotional Learning and Memory. Learning & Memory 7:257–266.

Holmes NM, Parkes SL, Killcross AS, Westbrook RF (2013) The basolateral amygdala is critical for learning about neutral stimuli in the presence of danger, and the perirhinal cortex is critical in the absence of danger. Journal of neuroscience 33:13112–13125.

Holmes NM, Cai SY, Lay BPP, Watts NR, Westbrook RF (2014) Extinguished second-order conditioned fear responses are renewed but not reinstated. Journal of Experimental Psychology: Animal Learning and Cognition 40:440.

Holmes NM, Fam JP, Clemens KJ, Laurent V, Westbrook RF (2022) The neural substrates of higher-order conditioning: a review. Neuroscience & Biobehavioral Reviews:104687.

Johansen JP, Cain CK, Ostroff LE, LeDoux JE (2011) Molecular mechanisms of fear learning and memory. Cell 147:509–524.

Kochli DE, Thompson EC, Fricke EA, Postle AF, Quinn JJ (2015) The amygdala is critical for trace, delay, and contextual fear conditioning. Learning & memory 22:92–100.

Lay BPP, Westbrook RF, Glanzman DL, Holmes NM (2018) Commonalities and differences in the substrates underlying consolidation of first- and second-order conditioned fear. The Journal of Neuroscience:2966–2917.

Leidl DM, Lay BPP, Chakouch C, Westbrook RF, Holmes NM (2018) Protein synthesis in the basolateral amygdala complex is required for consolidation of a first-order fear memory, but not for consolidation of a higher-order fear memory. Neurobiology of Learning and Memory 153:153–165.

Maren S, Ferrario CR, Corcoran KA, Desmond TJ, Frey KA (2003) Protein synthesis in the amygdala, but not the auditory thalamus, is required for consolidation of Pavlovian fear conditioning in rats. European Journal of Neuroscience 18:3080–3088.

Ostroff LE, Cain CK, Bedont J, Monfils MH, LeDoux JE (2010) Fear and safety learning differentially affect synapse size and dendritic translation in the lateral amygdala. Proceedings of the National Academy of Sciences 107:9418–9423.

Parkes SL, Westbrook RF (2010) The basolateral amygdala is critical for the acquisition and extinction of associations between a neutral stimulus and a learned danger signal but not between two neutral stimuli. Journal of Neuroscience 30:12608–12618.

Pavlov IP (1927) Conditioned reflexes (G. V. Anrep, Trans.).: London, England: Oxford University Press.

Paxinos G, Watson, C (2007) The rat brain in stereotaxic coordinates. Burlington: Academic.

Redondo RL, Morris RG (2011) Making memories last: the synaptic tagging and capture hypothesis. Nature Reviews Neuroscience 12:17–30.

Rescorla, R. A. (1973a). Effects of US habituation following conditioning. Journal of Comparative and physiological psychology 82:137.

Rescorla, R. A. (1973b). Second-order conditioning: Implications for theories of learning. In F. J. McGuigan & D. B. Lumsden, Contemporary approaches to conditioning and learning. V. H. Winston & Sons.

Rescorla RA (1982) Simultaneous second-order conditioning produces SS learning in conditioned suppression. Journal of Experimental psychology: animal Behavior processes 8:23.

Rizley RC, Rescorla RA (1972) Associations in second-order conditioning and sensory preconditioning. Journal of comparative and physiological psychology 81:1.

Schafe GE, LeDoux JE (2000) Memory consolidation of auditory pavlovian fear conditioning requires protein synthesis and protein kinase A in the amygdala. Journal of Neuroscience 20:RC96–RC96.

Schneider-Poetsch T, Ju J, Eyler DE, Dang Y, Bhat S, Merrick WC, Green R, Shen B, Liu JO (2010) Inhibition of eukaryotic translation elongation by cycloheximide and lactimidomycin. Nature chemical biology 6:209–217.

Tovote P, Fadok JP, Lüthi A (2015) Neuronal circuits for fear and anxiety. Nature Reviews Neuroscience 16:317–331.

Walker P, Carrive P (2003) Role of ventrolateral periaqueductal gray neurons in the behavioral and cardiovascular responses to contextual conditioned fear and poststress recovery. Neuroscience 116:897–912.

Witnauer JE, Miller RR (2011) Some determinants of second-order conditioning. Learning & Behavior 39:12–26.

